# TATA box-based analysis reveals activation of growth programs in bladder cancer cells despite ferroptosis activation

**DOI:** 10.1101/2025.06.05.658066

**Authors:** Mauricio Fernández-González, Ricardo Armisen, Mario I Fernández

## Abstract

A poorly studied mechanism suggests that growth is characterized by the expression of TATA-less genes under favorable conditions, while stress induces TATA-containing genes involved in stress responses. The role of this mechanism in cancer remains unclear. This study analyzes the presence of TATA boxes in gene expression data from bladder cancer (BC) cells undergoing ferroptosis (siUSP52 cells), a form of programmed cell death. We hypothesize that growth-related genes (TATA-less) are underrepresented in siUSP52 cells. We found a high prevalence of TATA-less genes in siUSP52 cells (62 of 90 differentially expressed), suggesting that BC promotes growth despite ferroptosis activation.

## Introduction

Bladder cancer (BC) is among the ten most common cancer malignancies, with an incidence of 613,791 new cases and a global mortality burden of 220,349 deaths in 2022, according to GLOBOCAN 2022 (Bray et al. 2024). BC exhibit aggressive behavior, metastatic potential, and limited treatment responsiveness (Sanli et al. 2017). With a 50% recurrence rate leading to poor outcomes (van Hoogstraten et al. 2023), further research is needed to uncover the molecular mechanisms driving BC progression.

A key point in the transition from normal to tumor cells is that cellular growth is driven by the upregulation of growth-related genes, primarily regulated by the mTOR- and MAPK-related signaling pathways (López-Maury, Marguerat and Bähler 2008). Under stress, MAPK-related pathways are activated, promoting the transcription of stress-response genes, which typically contain one or more TATA box motifs within their promoter regions. This activation simultaneously inhibits the transcription of growth-related genes that lack this motif. Conversely, under favorable conditions, mTOR signaling becomes active, promoting the transcription of growth-related genes that lack TATA box sequences and suppressing the expression of stress-response genes (López-Maury, Marguerat and Bähler 2008). This regulatory mechanism can be interpreted as a cellular strategy that enables the prioritization of resource allocation toward survival during stress and proliferation under optimal conditions (Fernández 2022;López-Maury, Marguerat and Bähler 2008).

TATA boxes are found in approximately 20% of eukaryotic genes (Guenther et al. 2007). The presence of TATA boxes is associated with increased transcriptional susceptibility (López-Maury et al. 2008; Muse et al. 2007; Radonjic et al. 2005) and noisy gene expression (López-Maury et al. 2008). Such variability and responsiveness are essential for cellular adaptation to stress in changing environments (Landry et al. 2007; López-Maury et al. 2008), enabling rapid and dynamic transcriptional regulation. In contrast, growth-related genes generally lack TATA boxes and therefore show lower transcriptional susceptibility, typically maintaining stable expression levels (López-Maury et al. 2008). This distinction highlights the TATA box as a key promoter element that helps differentiate stress-response genes from growth-related genes. Although the TATA box may play important roles in both stress adaptation and cell proliferation -and potentially influence cancer progression and persistence-its function in cancer remains largely unexplored.

In this study, we analyzed RNA-Seq transcriptional data from bladder cancer cells in which the USP52 gene was silenced (siUSP52). Silencing USP52 promotes ferroptosis, an iron-dependent form of cell death known for its tumor-suppressive effects (Stockwell et al. 2017). Since ferroptosis induces cellular stress, we hypothesize that growth-related genes, which typically lack TATA boxes, are underrepresented in siUSP52 cells, while stress-related genes, often containing TATA boxes, are overrepresented. Furthermore, since tumor cells can switch to alternative ways of producing energy under stress, such as the Warburg effect (WE), a metabolic shift that generates extra energy through glycolysis (Liberti and Locasale 2016), we hypothesize that siUSP52 cells might use this mechanism to keep growing despite ferroptosis. To test this, we analyzed the expression of glycolysis-related genes in siUSP52 cells to see if the Warburg effect is activated.

Our study offers a framework to better understand how the presence or absence of TATA box motifs affects stress response and growth in cancer cells, even when they are under stress. It also proposes a new way to classify gene functions based on whether they contain this motif. Additionally, understanding the role of TATA boxes may improve predictions of tumor behavior and treatment responses, especially for therapies that induce cellular stress.

## Results

We identified 90 differentially expressed genes, of which 69 were overexpressed. Among these overexpressed genes, 18 contained TATA boxes and 45 did not. Of the 21 underexpressed genes, 3 contained TATA boxes and 17 did not (see gene list in Supplementary Material S3). Therefore, in siUSP52 cells, growth-related genes (TATA-less genes) were expressed at higher levels than stress-related genes (TATA-containing genes) (Fig. 1). Additionally, the fold-change analysis showed that growth-related genes had larger changes in expression than stress-related genes, with overexpression being more common than underexpression in both groups (Fig. 2). Interestingly, although fewer stress-related genes were expressed and their changes in expression were not greater than those of growth-related genes, the stress-related genes showed greater variability in expression compared to growth-related genes (Fig. 2). This finding aligns with previous reports indicating that genes rich in TATA boxes tend to exhibit more variable, or “noisy,” transcriptional activity (López-Maury, Marguerat, and Bähler 2008), and it also serves as a confirmatory analysis to validate the functional assignment of genes to both biological categories based on TATA box identification.

**Figure 1.**
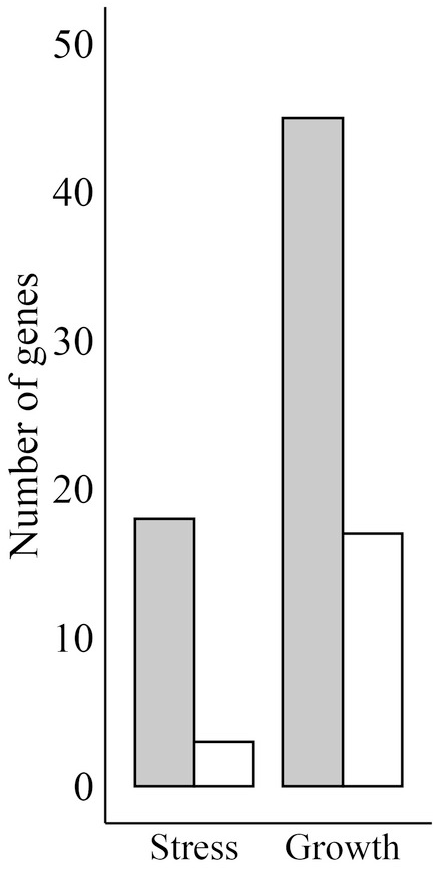
Number of over and underexpressed genes related to stress (TATA-containing) and growth (TATA-less) in in siUSP52. Gray bars represent the number of overexpressed genes, and white bars represent the number of underexpressed genes.

**Figure 2.**
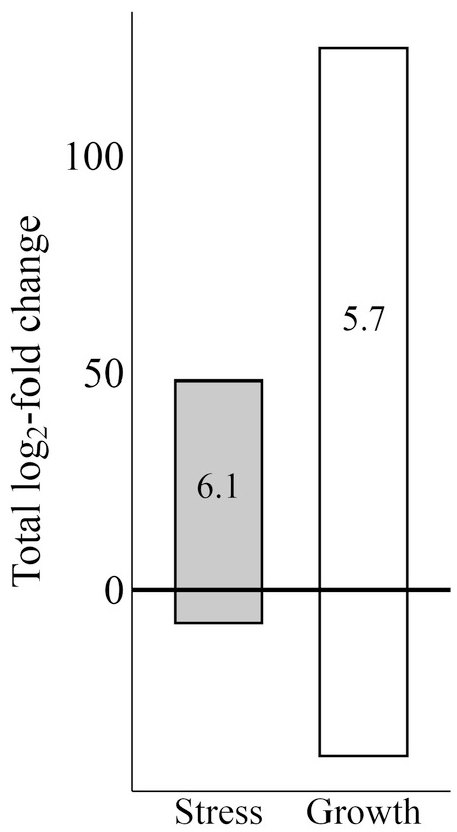
Gene expression of stress genes (TATA-containing) and growth genes (TATA-containing) in siUSP52. The Y-axis shows the sum of log2 fold changes. The zero line separates overexpression (sum of positive values) from underexpression (sum of negative values). Numbers inside the bars indicate the standard deviation of the fold-change values.

The higher abundance of TATA-containing genes and the low expression of stress-related genes suggest that siUSP52 cells are promoting cellular growth despite the activation of ferroptosis. Although this initially suggests that the cells are not under stress -since growth does not appear to be inhibited by MAPK activation-the expression of WE-like genes (Table 1) suggests that the cells may be receiving an additional energy input, which could prevent the activation of MAPK-related pathways and help sustain active mTOR signaling, thereby promoting the transcription of growth-related genes. These findings do not support our initial hypothesis, which predicted that siUSP52 cells would exhibit reduced expression of growth-related genes. We propose that the critical factor underlying our results is the observed overexpression of glycolysis-related genes (Table 1), which likely provides the energy necessary to (i) maintain cell growth by sustaining mTOR signaling and preventing MAPK activation, and (ii) support stress response mechanisms, albeit to a lesser extent, to mitigate ferroptosis-induced damage. In the following sections, we explore both aspects to better understand how cells manage to sustain growth despite the activation of ferroptosis, and, given the observed noisy activation of stress-related genes, we also examine the potential role of TATA-containing gene overexpression as an adaptive mechanism for coping with stress conditions.

**Table 1.** Genes with increased expression involved in glycolysis in siUSP52 cells. The first column, “Gene,” lists the gene IDs. The “Glycolysis/Expression” column first indicates the effect of each gene’s expression on glycolysis (“Glycolysis”) and then whether the gene was found to be overexpressed or underexpressed in siUSP52 cells. “Reference” provides the corresponding bibliography for the glycolytic effect. Asterisks indicate TATA-containing genes involved in glycolysis activation.

| Gene | Glycolysis/Expression | Reference |
| --- | --- | --- |
| GDF15 | Promote/UP | Xue et al. 2025 |
| GPR65 | Promote/UP | D Patel pers. Comm. |
| PPARGC1A | Decrease/UP | Zuo et al. 2021 |
| PTGS2 * | Promote/UP | Zhang et al. 2018 |
| SPP1 | Promote/UP | Lu et al. 2020 |
| CA8 * | Promote/UP | Wang et al. 2016 |
| CFH | Promote/UP | Armento et al. 2020 |
| LIPH | Decrease/UP | Han et al. 2023 |
| MYCT1 * | Promote/UP | Ding et al. 2022 |
| NGF | Decrease/UP | Fodelianaki et al. 2019 |
| MYB | Promote/Down | Liu et al. 2022 |
| PIGX | Decrease/Down | Fineran et al. 2007 |
| TYMS | Promote/Down | Li et al. 2023 |

### Glycolysis and growth promotion in ferroptosis-activated cells

As previously mentioned, ferroptosis is an iron-dependent, non-apoptotic form of cell death with strong antitumor potential. It is triggered by oxidative damage to membrane phospholipids, resulting from free intracellular iron and lipid peroxidation (Yao et al. 2021). Despite its antitumor capacity, tumor cells may evade ferroptosis by upregulating glycolysis, which increases the production of NADPH and glutathione, both essential for the activity of GPX4, a key enzyme that reduces lipid peroxides and prevents ferroptosis activation (Yao et al. 2021). Therefore, glycolysis appears to play a dual role: it not only provides energy to support cell growth but also helps suppress ferroptosis, thereby maintaining cell viability and promoting continued proliferation.

### Variability in the expression of TATA-containing genes: an adaptive response to uncertainty

The higher variability exhibited by stress-related genes (Fig. 2) is attributable to the presence of TATA boxes and enables organisms to prepare not only for specific stress conditions but also for novel or unpredictable environmental challenges. This transcriptional noise plays a key role in enhancing survival under adverse conditions by promoting phenotypic diversity within the cell population, increasing the likelihood that some cells will withstand environmental stress (Barkai and Shilo 2007; Blake et al. 2006; Newman et al. 2006; Raser and O’Shea 2004; Acar, Mettetal and Van Oudenaarden 2008; López-Maury, Marguerat and Bähler 2008). However, its benefits have limits, because excessive variability may lead to a loss of response specificity and compromise the robustness of essential cellular functions (Barkai and Shilo 2007; Gerhart and Kirschner 2007; Lenski, Barrick and Ofria 2006; Williams and Stein 2004).

In cancer cells, the noisy expression conferred by TATA-containing genes can be beneficial for promoting survival and growth by increasing resistance not only to external stressors but also to internal changes. For example, our results (Table 1) show that some TATA-containing genes are associated with the promotion of glycolysis, potentially enabling cells to bypass ferroptosis while simultaneously providing the energy needed to sustain growth. This transcriptional noise may facilitate resistance to both current and emerging anticancer therapies, thereby reducing treatment efficacy. We hypothesize that even at low levels, the expression of TATA-containing genes is maintained to preserve the ability to respond to unpredictable conditions.

Based on our analysis of the TATA box motif, we conclude that siUSP52 cells exhibit a growth-oriented transcriptional profile, characterized by higher expression of growth-related genes and lower expression of stress-related genes, a result contrary to our initial hypothesis. This growth-oriented state may be sustained by the upregulation of glycolysis-related genes, as this metabolic pathway not only increases the energy available to support growth but also helps mitigate ferroptosis activation.

## Materials and Methods

We used the RNA-Seq expression data of two types of T24 bladder cancer cells siNC and siUSP52, available in Liu et al (2024) and in Gene Expression Omnibus (https://www.ncbi.nlm.nih.gov/geo/), reference number GSE263111. siNC are bladder cancer cells while siUSP52 are bladder cancer cells with silenced USP52 gene leading to activation of ferroptosis. This dataset consists of a table that includes the Gene ID, fold change, log2(fold change), p-value, q-value and gene expression data for all genes sequenced in both cell types. We utilized R programming language (R Core Team, 2024) to filter RNA-Seq data using a p-value < 0.001, q-value <0.001 and a log2(fold change) > 2 or ≤ –2 (R script is available as supplementary materials S1). Using the gene IDs of the filtered genes, we downloaded their nucleotide sequences in FASTA format from ENSEMBL, including 1000 base pairs upstream from the gene start to ensure that the promoter region was included. Next, we analyzed the presence of TATA boxes in the FASTA sequences. The main goal of this phase was not just to identify any TATA box-like sequences, since the true TATA boxes are located approximately 26 base pairs upstream of the transcription start site (TSS). To achieve this goal, we first identified the transcription start site (TSS) within each FASTA sequence by analyzing approximately the first 1500 base pairs using the BDGP web tool, which predicts TSS in eukaryotic sequences provided in FASTA format (https://www.fruitfly.org/seq_tools/promoter.html). After identifying the TSS for all gene sequences, we used the R language (R Core Team, 2024) to assess the presence of a TATA box in the vicinity of the TSS. We developed a script that searches within a region approximately 26 base pairs upstream for at least one of four consensus sequences for the TATA box: TATAAAA, TATAAAT, TATATAA, and TATATAT (R script is available as supplementary material S2). Finally, we obtained a table with two columns: the first containing the gene ID, and the second indicating whether a TATA box sequence was found near the TSS (‘Yes’ or ‘No’). As mentioned earlier, genes containing a TATA box motif were classified as stress-related (TATA-containing), whereas those lacking the motif were classified as growth-related (TATA-less).

To assess whether tumor cells with activated ferroptosis favor growth, stress responses, or both, we counted the number of growth- and stress-related genes that were overexpressed or underexpressed in siUSP52 cells. This quantification would indicate, for example, that a higher number of overexpressed genes in one group provides more options to promote the corresponding biological function. To assess the intensity of the transcriptional response, we compared the fold changes of stress-related and growth-related genes in siUSP52 cells. The difference between these values was used as an indicator of the relative transcriptional effort invested in promoting each biological function.

Since previous work has predicted that stress-related genes exhibit noisy transcription (López-Maury, Marguerat, and Bähler 2008), we measured the variability of the transcriptional response in siUSP52 cells by calculating the standard deviation of the fold-change for both stress- and growth-related genes. This calculation serves as a confirmatory analysis to validate the functional assignment of genes to the two biological categories based on expression variability.

Finally, to analyze energy availability in the dynamic balance between growth and stress response, and considering that some tumor cells can activate the Warburg effect to fuel biological functions, we evaluated the activation of glycolysis-related genes in siUSP52 cells. For this purpose, we identified each gene’s function by searching the gene ID in the GeneCards database (https://www.genecards.org/) and reviewed relevant scientific literature on the genes and glycolysis using PubMed (https://pubmed.ncbi.nlm.nih.gov/).

## Acknowledgments

During the preparation of this work, the authors used ChatGPT-4 in order to assist with language refinement and text structuring. After using this tool, the authors reviewed and edited the content as needed and takes full responsibility for the content of the published article. The authors have reviewed and edited the output and take full responsibility for the content of this publication. This research was funded by Agencia Nacional de Investigación y Desarrollo, under the grant Proyecto Anillo ACT210079.

## Author Contributions

Conceptualization, M.I.F. and M.F.; methodology, M.F.; software, M.F.; formal analysis, M.F.; investigation, M.I.F., R.A. and M.F.; resources, M.I.F. and R.A.; data curation, M.F.; writing-original draft preparation, M.F.; writing-review and editing, M.I.F., R.A. and M.F.; visualization, M.F.; supervision, M.I.F. and R.A; project administration, M.I.F. and R.A.; funding acquisition, M.I.F. and R.A. All authors have read and agreed to the published version of the manuscript.

## Supplementary Materials

The list with TATA-containing and TATA-less genes and informatic script are available as supplementary materials.

